# Redundant *SCARECROW* genes pattern distinct cell layers in roots and leaves of maize

**DOI:** 10.1101/567156

**Authors:** Thomas E. Hughes, Olga V. Sedelnikova, Hao Wu, Philip W. Becraft, Jane A. Langdale

**Affiliations:** Department of Plant Sciences, University of Oxford, South Parks Road, Oxford, UK, OX1 3RB; Genetics, Development, and Cell Biology Department, Iowa State University, Ames, Iowa, USA, 50011; Current address: Syngenta Jealott’s Hill International Research Centre, Bracknell RG42 6EY

**Keywords:** Radial-patterning, SCARECROW, Kranz, Mesophyll, Bundle-sheath, Maize

## Abstract

The highly efficient C4 photosynthetic pathway is facilitated by ‘Kranz’ leaf anatomy. In Kranz leaves, closely spaced veins are encircled by concentric layers of photosynthetic bundle sheath (inner) and mesophyll (outer) cells. Here we demonstrate that in the C4 monocot maize, Kranz patterning is regulated by redundant function of SCARECROW 1 (ZmSCR1) and a previously uncharacterized homeolog ZmSCR1h. *ZmSCR1* and *ZmSCR1h* transcripts accumulate in ground meristem cells of developing leaf primordia and in *Zmscr1;Zmscr1h* mutant leaves, most veins are separated by one rather than two mesophyll cells; many veins have sclerenchyma above and/or below instead of mesophyll cells; and supernumerary bundle sheath cells develop. The mutant defects are unified by compromised mesophyll cell development. In addition to Kranz defects, *Zmscr1;Zmscr1h* mutants fail to form an organized endodermal layer in the root. Collectively, these data indicate that ZmSCR1 and ZmSCR1h redundantly regulate cell-type patterning in both leaves and roots of maize. Leaf and root pathways are distinguished, however, by the cell layer in which they operate – mesophyll at a two-cell distance from leaf veins versus endodermis immediately adjacent to root vasculature.

**Summary statement:** Two duplicated maize *SCARECROW* genes control the development of the endodermis in roots and the mesophyll in leaves

## INTRODUCTION

The C4 photosynthetic pathway, which is responsible for around 21% of global primary productivity despite being found in only ~3% of plant species (Ehleringer et al., 1997; Sage et al., 2011), is underpinned by a specialized leaf anatomy known as Kranz (the German word for wreath) (reviewed in Sedelnikova et al., 2018). Unlike in C3 plants, where photosynthesis only occurs in the mesophyll cells, the C4 pathway is separated between bundle sheath (BS) and mesophyll (M) cells, with the two cell-types forming concentric wreaths around leaf veins (reviewed in Langdale, 2011). Efficient operation of the C4 cycle relies on an increased BS to M cell ratio relative to that seen in C3 plants, an increase that is achieved by altering vein density so that vascular bundles are often separated by only two mesophyll cells in a recurring vein-BS-M-M-BS-vein pattern across the leaf. In C4 monocots such as maize, this high vein density results from the formation of small ‘rank-2’ intermediate veins in between the lateral and rank-1 intermediate veins that are common to both C3 and C4 species (Esau, 1943; Russell and Evert, 1985; Sharman, 1942). Given the higher yields found in many C4 plants, there are ongoing attempts to engineer the C4 pathway into C3 crops (Hibberd et al., 2008; von Caemmerer et al., 2012; Wang et al., 2016), however, such attempts require a far better understanding of how vein spacing and leaf cell fate is regulated in C4 species.

Our understanding of the genetic components that regulate the development of Kranz anatomy is extremely limited, in part because traditional approaches to gene-discovery, such as mutant screens, failed to reveal any regulators of vein spacing or BS/M cell fate (Langdale, 2011). More recent transcriptomic analyses identified candidate genes that are expressed in a manner consistent with roles in Kranz patterning (Fouracre et al., 2014; Wang et al., 2013), but in most cases gene function has yet to be validated. One candidate, the maize GRAS protein SCARECROW1 (ZmSCR1), has been shown to regulate aspects of Kranz patterning in that *Zmscr1* mutants have subtle alterations in vascular, BS and M development (Slewinski et al., 2012). In *Arabidopsis thaliana* (hereafter referred to as Arabidopsis) the *ZmSCR1* ortholog radially patterns cell-types in the root (Di Laurenzio et al., 1996; Wysocka-Diller et al., 2000); AtSCR prevents movement of AtSHORTROOT (AtSHR) beyond the cell-layer adjacent to the vasculature, which ensures specification of endodermal cells in that layer (Cui et al., 2007). However, an organized endodermal cell layer is present in *Zmscr1* mutants (Slewinski et al., 2012), suggesting that gene function may have diverged between maize and Arabidopsis. Given that the root endodermis and the leaf BS are considered analogous cell types (Esau, 1943; Nelson, 2011), it is possible that an ancestral SCR patterning function was recruited in the leaf rather than the root in maize, but the subtle phenotype reported in leaves of *Zmscr1* mutants precludes an understanding of the precise role played during Kranz development.

Both gene and whole genome duplication events are highly prevalent throughout the plant phylogeny (Adams and Wendel, 2005; Blanc and Wolfe, 2004) and if retained in the genome, duplicated genes are free to sub- or neo-functionalize (Moore and Purugganan, 2005; Ohno, 1970). Perhaps more commonly, however, gene duplicates function redundantly. Indeed, there are many examples illustrating the importance of genetic redundancy in plants, and without understanding phylogenetic context, loss-of-function data can be difficult to interpret (Strable et al., 2017; Yi et al., 2015). This is particularly important in maize, which in addition to undergoing three ancient whole genome duplication events common to monocots, has also undergone a more recent event not shared with its close relative *Sorghum bicolor* (Messing et al., 2004; Schnable et al., 2009; Swigonova et al., 2004). It is thus likely that *ZmSCR1* acts redundantly with a duplicate gene to pattern cell-types in maize.

To better understand the role of ZmSCR1 in maize development we first constructed a phylogeny of SCR-related genes, which revealed that *ZmSCR1* has a previously overlooked homeolog duplicate *ZmSCR1h*. When transposon-induced loss of function alleles of both genes were combined, double mutants exhibited leaf and root phenotypes that were not seen in segregating single mutant siblings. Cell-type specification was perturbed in both the leaf and root of *Zmscr1; Zmscr1h* double mutants, with endodermal defects observed in the root. Intriguingly, however, M rather than BS cell development was primarily perturbed in the leaf. We present a quantitative analysis of single and double *Zmscr1; Zmscr1h* mutant leaf phenotypes, plus expression data for both genes in developing wild-type maize leaf primordia. The results are discussed in the context of how SCR function has diversified in flowering plants.

## RESULTS

### *SCR* is duplicated in maize

To determine phylogenetic relationships between SCR-related genes in land plants, a maximum likelihood phylogeny was constructed. Figure 1A shows that two clades of *SCR* genes are present in both eudicots and monocots, with the underlying duplication event inferred after the divergence of *Physcomitrella patens* and vascular plants. In Arabidopsis, the *SCR* clade contains a single gene (*AtSCR*), with the closest related homolog (*SCR-LIKE 23 - AtSCL23*) in the sister clade. Apart from *Ananas comosus*, the sampled monocot genomes contain single-copy orthologs of *AtSCL23*. In contrast, *SCR* has independently duplicated in at least four monocot genomes (maize, *Sorghum bicolor, Setaria italica* and *Oryza sativa*), and the apparent single copy in *Setaria viridis* is likely an annotation error. The maize *SCR* duplicates reside on syntenic regions of chromosomes 4 (*ZmSCR1*) and 2 (*ZmSCR1h*), and have previously been annotated as likely homeolog gene pairs that arose through the recent maize whole genome duplication (Schnable et al., 2011). Sequence comparisons reveal 85% amino acid identity between ZmSCR1 and ZmSCR1h and both contain an N-terminal domain that prevents intercellular movement of the AtSCR protein (Gallagher and Benfey, 2009). These observations suggest that ZmSCR1 and ZmSCR1h act redundantly, and in a cell-autonomous manner.

**Figure 1.**
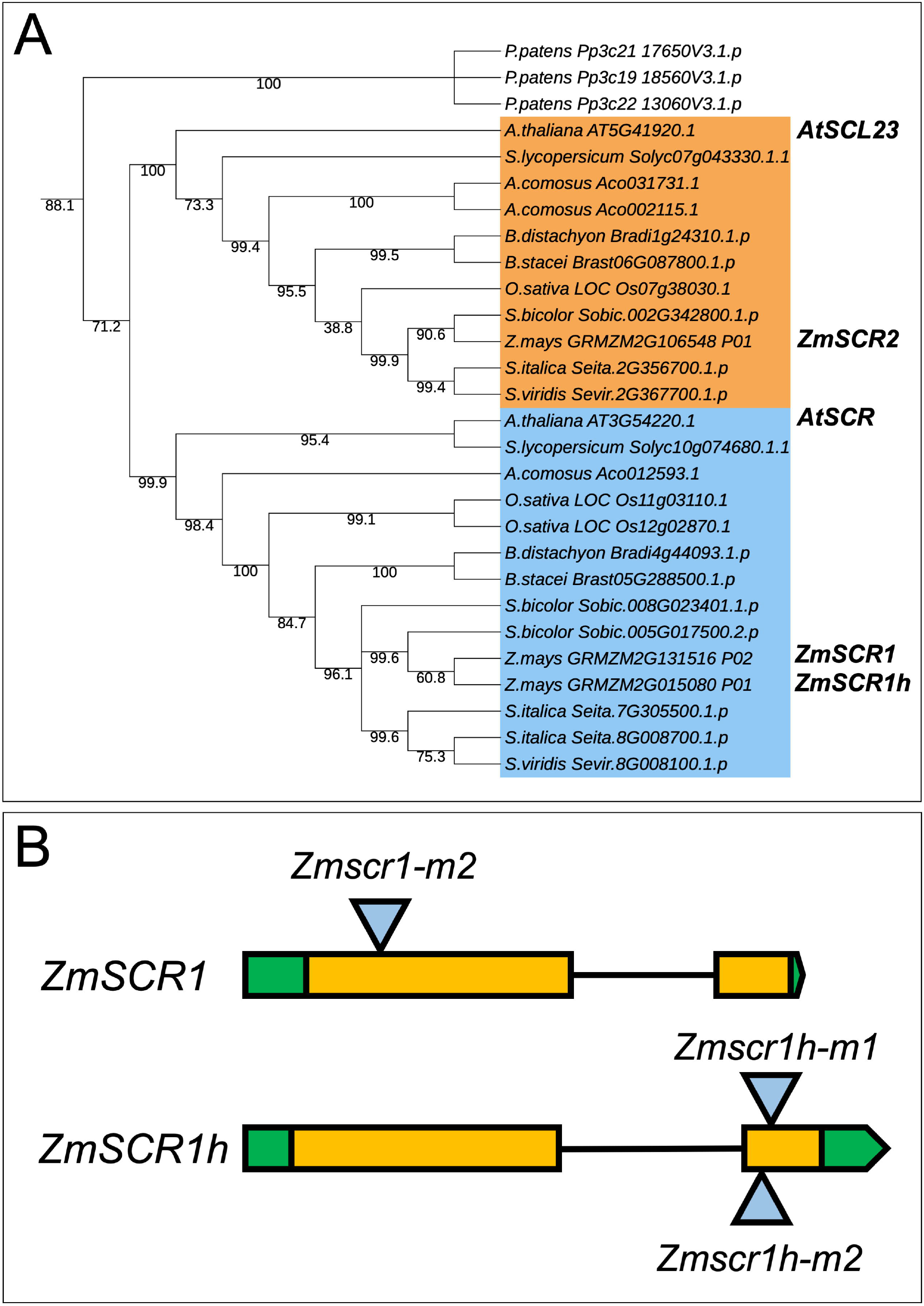
Transposon insertions in maize *AtSCR* orthologs. **A)** Maximum likelihood phylogeny of *SCR* genes. Bootstrap values are indicated below branches. Light blue shading indicates the *AtSCR* clade, light orange shading indicates the *AtSCL23* clade. *Physcomitrella patens* sequences were included as an outgroup. **B)** Cartoon depiction of *Mutator* transposon insertions in *ZmSCR1* and *ZmSCR1h*. All three alleles were in the W22 inbred background from the UniformMu project. UTRs (green), exons (orange), introns (black line) and transposon insertion site (blue triangle) are indicated.

### Transposon insertion alleles of *ZmSCR1* and *ZmSCR1h* cause loss of function

To test the hypothesis of functional redundancy, we first identified transposon insertion alleles for each gene. Two *Zmscr1* alleles (*-m1* and *-m2*) have been reported previously (Slewinski et al., 2012), and we identified two independent *Zmscr1h* alleles (*Zmscr1h-m1* and *-m2*) in the UniformMu transposon insertion collection (Fig. S1A) (McCarty et al., 2005). In both *Zmscr1h-m1* and *-m2*, a *Mutator* (*Mu*) element was predicted to be inserted in the second exon of the *ZmSCR1h* coding sequence (Fig. 1B). Seed stocks for both alleles, plus *Zmscr1-m2* which is in the same UniformMu W22 background, were obtained from the Maize Genetics Stock Centre (http://maizecoop.cropsci.uiuc.edu/). Single *Mu* insertions in the genes of interest are documented for the *Zmscr1-m2* and *Zmscr1h-m1* lines, whereas the *Zmscr1h-m2* line contains four additional *Mu* elements inserted at other loci (Fig. S1A). Insertion positions were confirmed by polymerase chain reaction (PCR) amplification of genomic DNA, using primers in the transposon and in the adjacent genic region (Fig. S1B, C). In all cases, the size of the amplified product was consistent with the predicted insertion site. Primers flanking the *Mu* element enabled homozygous mutant individuals to be identified.

To confirm that the transposon insertion alleles compromised gene function, transcripts were amplified and sequenced, using RNA extracted from homozygous mutant leaf primordia as a starting template. Reverse transcriptase (RT)-PCR revealed that in all cases, the *Mu* element was present in the *ZmSCR1* or *ZmSCR1h* transcript, at the position predicted by the insertion site (Fig. S1D). As such, even if transcripts were translated, a non-functional protein would be produced.

### Loss of function *ZmSCR1h* mutants do not exhibit cell-type patterning defects

To determine whether *Zmscr1h* mutants display similar defects in Kranz patterning to those reported in *Zmscr1* mutants (Slewinski et al., 2012), leaf traits were compared between *Zmscr1-m2, Zmscr1h-m1* and *Zmscr1h-m2* single mutants, and corresponding wild-type siblings segregating in each line. Figure 2 shows that there was no qualitative difference between wild-type and either *Zmscr1* or *Zmscr1h* single mutants in overall plant growth (Fig. 2A-D), or in general Kranz patterning (Fig. 2E-H). Quantification of the number of M cells between veins (Fig. 2I), vein density across the leaf (Fig. 2J), and the ratio of rank-1:rank-2 intermediate veins (Fig. 2J), failed to confirm previous reports of altered vein density and interveinal M cell number in *Zmscr1-m2* mutants, but showed that *Zmscr1-m2* mutants exhibit a small but significant increase in the ratio of rank-1:rank-2 intermediate veins (Fig. 2J). No significant difference was observed between wild-type and either *Zmscr1h* mutant allele for any of the measured traits. Previous reports of supernumerary BS cells in *Zmscr1-m2* mutants (Slewinski et al., 2012) were confirmed, and the trait was also seen in *Zmscr1h-m1* and *Zmscr1h-m2* single mutants (Fig. 2K). However, very few instances were observed above background levels in segregating wild-type siblings and over 97% of veins in mutant leaves were surrounded by a normal BS cell layer (Fig. 2K). As such, it can be concluded that mutations in *ZmSCR1* or *ZmSCR1h*cause only minor perturbations to Kranz patterning mechanisms.

**Figure 2.**
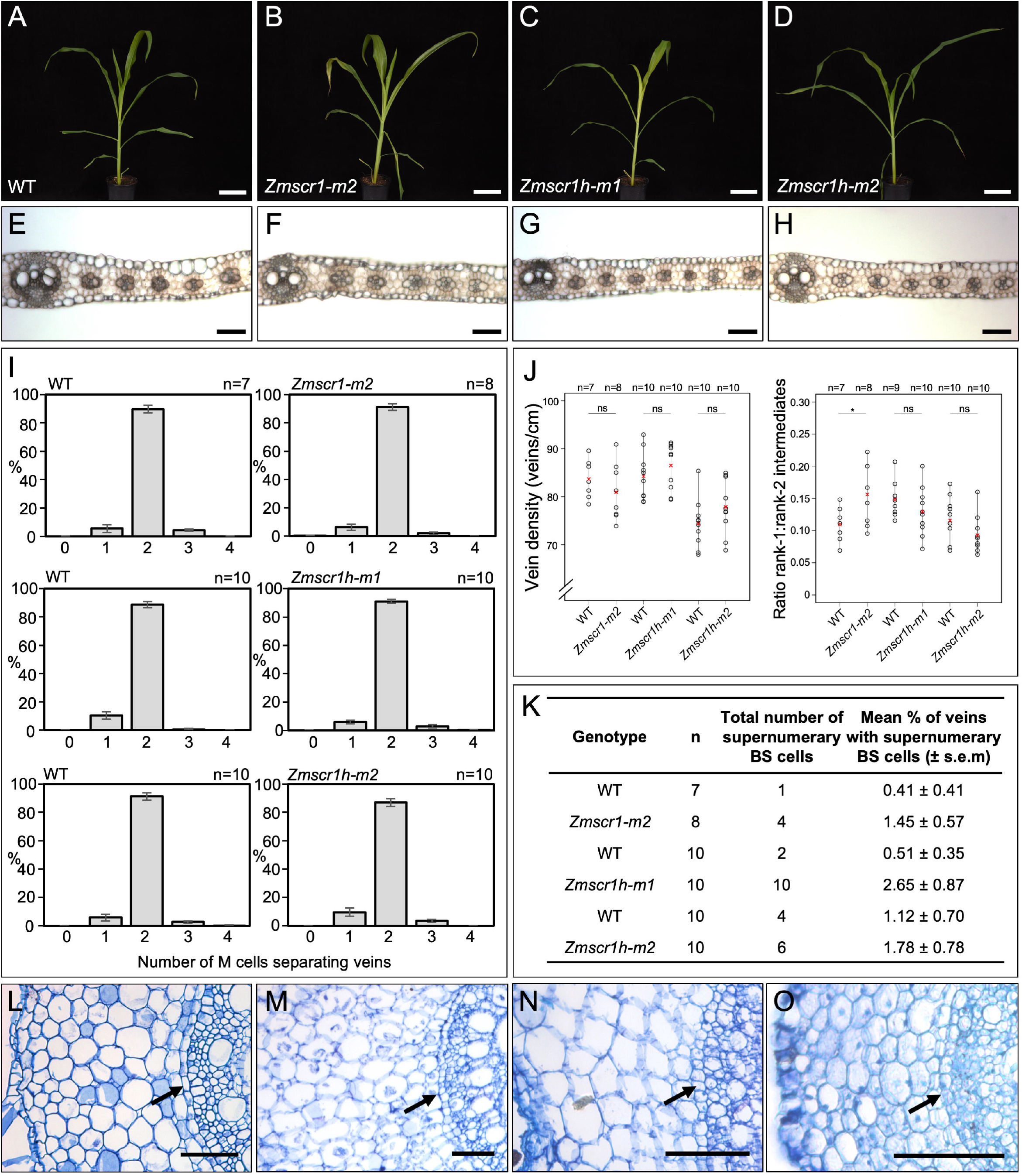
Phenotype of *Zmscr1* and *Zmscr1h* single mutants. **A-D)** Representative whole plant phenotype at 32 days after planting. **E-H)** Representative transverse sections of fully expanded leaf 5 of WT (E), *Zmscr1-m2* (F) *Zmscr1h-m1* (G) and *Zmscr1h-m2* (H) plants. **I)** Quantification of mean % of M cells between vein pairs. In each case, the WT plot on the left represents a segregating sibling from the same family as the mutant presented in the corresponding plot on the right. Error bars are standard errors of the mean. **J)** Quantification of vein density and the ratio of rank1 to rank 2 intermediate veins in leaf 5. In each case, data from segregating WT (left) and corresponding mutant (right) are presented. Means are indicated by red crosses. Statistical significance between WT and mutant was assessed using Student’s t-tests (two-tailed): ns = no significant difference; * indicates p ≤ 0.05. **K)** Quantification of mean number of supernumerary BS cells in leaf 5. In each case, data from segregating WT (top) and corresponding mutant (bottom) are presented. s.e.m indicates standard error of the mean. **L-O)** Representative transverse sections of seminal roots of WT (L), *Zmscr1-m2* (M) *Zmscr1h-m1* (N) and *Zmscr1h-m2* (O) plants. Arrows indicate the endodermal layer positioned between vasculature and cortex. Scale bars = 10cm (A-D); 100μm (E-H & L-O)

In the absence of any major defects in *Zmscr1h* mutant leaves, and given that root development is perturbed in Arabidopsis mutants, we also examined cell-type patterning in single mutant roots. In *Atscr* mutants, instead of distinct layers of endodermis and cortex differentiating around the vasculature, a single cell layer with characteristics of both cell-types forms (Di Laurenzio et al., 1996). Notably, qualitative histological analysis of *Zmscr1-m2, Zmscr1h-m1* and *Zmscrh1-m2* root sections revealed normal development, with a clear endodermal boundary between the vasculature and the multiple cortical layers that are characteristic in maize (Fig. 2L-O). This observation suggests either that *ZmSCR1* and *ZmSCR1h* act redundantly to form an endodermal layer in the maize root, or that the role of the SCR pathway in patterning endodermis has diverged between Arabidopsis and maize.

### *Zmscr1;Zmscr1h* double mutants exhibit stunted growth

To distinguish hypotheses of redundant versus divergent function of ZmSCR1 and ZmSCR1h, double mutant lines were generated with *Zmscr1-m2* and the two independent *Zmscr1h* alleles. In F2 populations, 6/107 and 5/78 individuals were genotyped as *Zmscr1-m2;Zmscr1h-m1* and *Zmscr1-m2;Zmscr1h-m2* respectively, which matched the expected segregation ratio (χ^2^ p>0.05 in both cases). Although *Zmscr1;Zmscr1h* double mutants sometimes formed tassels when grown in the greenhouse, they rarely produced ears, and plants were never successfully self-pollinated. As such, phenotypic analysis was initially carried out using F2 populations segregating 1 in 16 for the homozygous double mutants. More detailed characterization was undertaken with F3 progeny of self-pollinated F2 *Zmscr1-m2/+;Zmscr1h-m1* plants, which segregated 1 in 4 for the *Zmscr1-m2;Zmscr1h-m1* homozygous double mutants. In all experiments, comparisons were made to wild-type plants segregating in the same population.

Unlike segregating single mutants, double mutants exhibited slower growth and reduced plant size (Fig. 3), similar to that reported for *Atscr* mutants (Di Laurenzio et al., 1996). Furthermore, *Zmscr1-m2;Zmscr1h-m1* and *Zmscr1-m2;Zmscr1h-m2* mutant leaves were on average 46% and 74% the length of corresponding wild-type leaves respectively, with leaf width also proportionally reduced (Fig. S2A, B). Despite the plants being smaller, they were not markedly developmentally delayed, because the emergent leaf in *Zmscr1-m2;Zmscr1-m1* double mutants was only around one plastochron (the time interval between initiation of leaves at the shoot apex) behind that of wild-type after 31 days of growth (Fig. S2D). Strikingly, however, emerging leaves in both double mutants were droopy (Fig. 3C, D). This phenotype was associated with a reduction in midrib length such that it extended on average 41% (*Zmscr1-m2;Zmscr1h-m1*) and 61%(*Zmscr1-m2;Zmscr1h-m2*) of the total leaf length compared to 75% and 83% in leaves of wild-type siblings (Fig. S2C). Defective growth was more severe in field-grown plants, suggesting an environmental influence.

**Figure 3.**
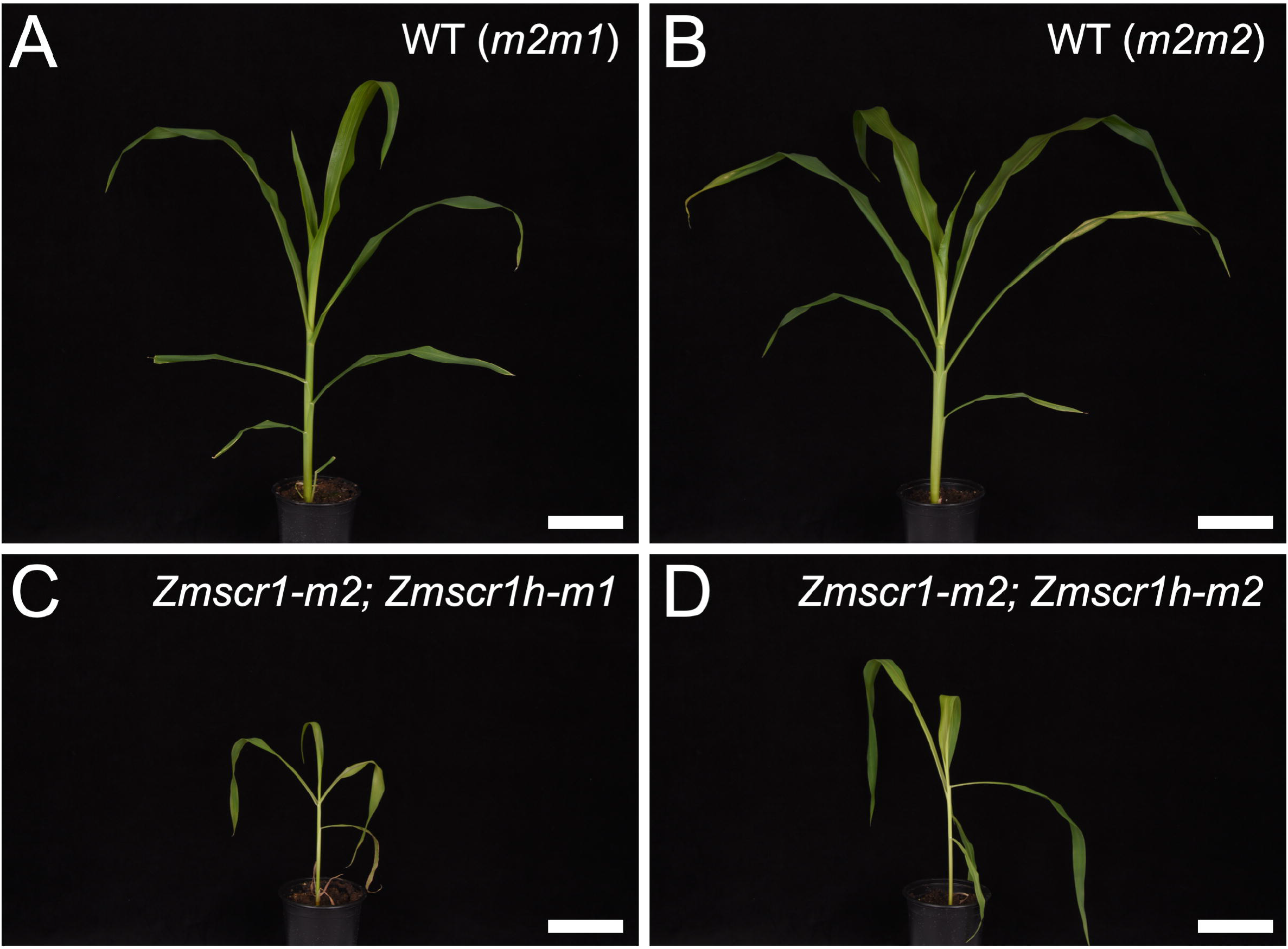
Growth is stunted in *Zmscr1;Zmscr1h* mutants. **A-D)** 32 day old segregating WT (A,B) and *Zmscr1;Zmscr1h* double mutant siblings (C,D). Scale bars = 10cm.

### ZmSCR1 and ZmSCR1h redundantly regulate formation of the endodermis in maize roots

Given the environmental influence on the stunted growth phenotype of *Zmscr1;Zmscr1h* double mutants, and the well-characterized phenotype of *Atscr* mutants, we hypothesized that loss of *ZmSCR1* and *ZmSCR1h* function disrupted differentiation of the root endodermis. Transverse sections from the maturation zone of *Zmscr1-m2;Zmscr1h-m1* primary roots were therefore examined. Figure 4 shows that overall size and structure of the root is normal, with apparently typical differentiation of cortex cells (Fig. 4A, B). However, there is no clear boundary between the vasculature and cortex (Fig. 4E). This pattern was also apparent in the seminal roots of *Zmscr1-m2;Zmscr1h-m2* mutants (Fig. 4C, F). These results demonstrate that ZmSCR1 and ZmSCR1h redundantly regulate formation of an organized endodermal layer in maize roots, possibly in an analogous manner to AtSCR in Arabidopsis.

**Figure 4.**
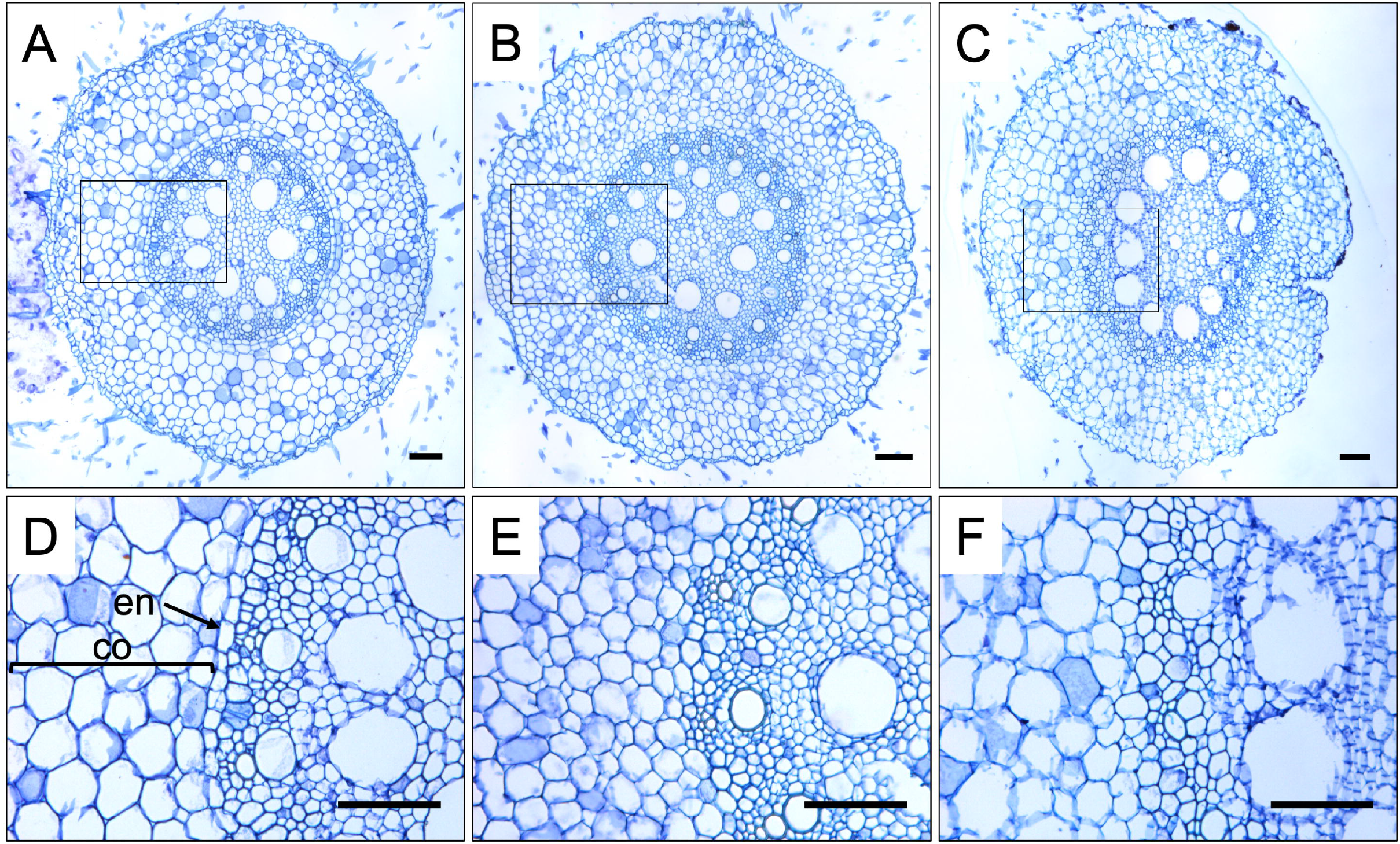
ZmSCR1 and ZmSCR1h regulate endodermis formation during root development. Representative transverse sections of roots from WT plants segregating in the *Zmscr1-m2;Zmscr1h-m1* background (A, D) plus *Zmscr1-m2;Zmscr1h-m1* (B, E) and *Zmscr1-m2;Zmscr1h-m2* (C, F) double mutants. Sections were taken from the maturation zone of either primary roots 4 days after germination (A, B, D, E) or seminal roots 35 days after germination (C, F). D, E and F are enlargements of the area indicated by the rectangle in the corresponding whole root image in the panel above. Cortex (co) and endodermis (en) are indicated. Scale bars = 100μm.

### Bundle sheath cell specification is not perturbed in *Zmscr1;Zmscr1h* mutant leaves

The original thesis that the SCR pathway may regulate Kranz patterning in the maize leaf was predicated on the long held view that the root endodermis and leaf BS are analogous cell-types, and that radial patterning mechanisms may be conserved in root and leaf (Slewinski, 2013). Given the clear absence of an organized endodermal layer in *Zmscr1;Zmscr1h* double mutants (Fig. 4), if leaf and root pathways are conserved, BS cell formation should be severely perturbed in leaves. To test whether this is the case, the position and number of BS cells was quantified across double mutant leaves (Fig. 5A-C). Crucially, all veins had a normal BS cell layer (Fig. 5A, B) and as such, whereas the SCR pathway is necessary for the formation of an endodermal layer in the root, it is not required for the development of a BS cell layer around leaf veins.

**Figure 5.**
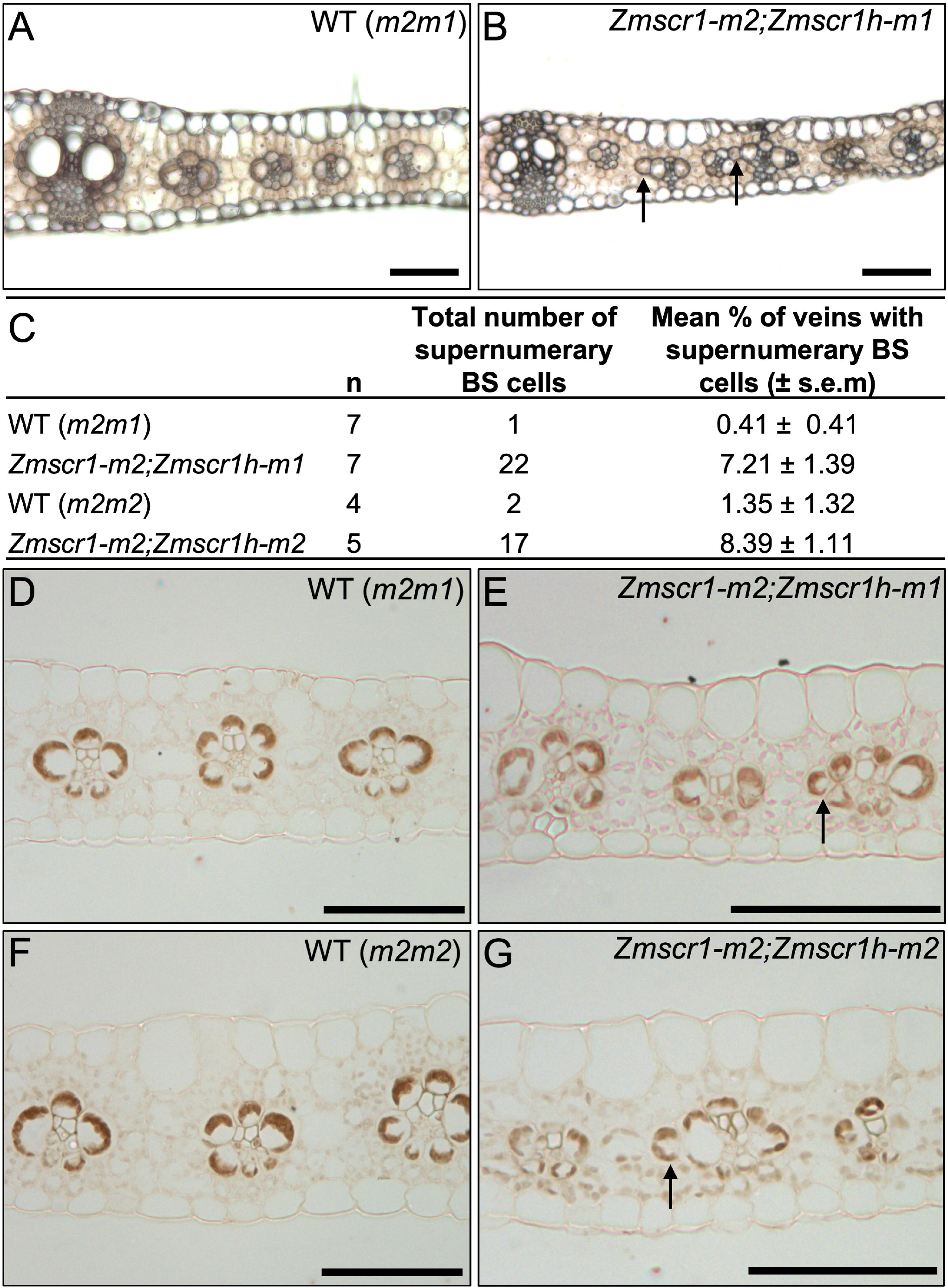
*Zmscr1;Zmscr1h* mutants form supernumerary BS cells. **A-B)** Representative fresh cut transverse sections of fully expanded leaf 5 from WT (*m2m1* segregant) (A) and *Zmscr1-m2;Zmscr1h-m1* mutant (B) plants. **C)** Quantification of the frequency of extra BS cells in different mutant backgrounds. s.e.m indicates standard error of the mean. WT (*m2m1*) data are also presented in Fig. 2K. **D-G)** Immunolocalization of ZmNADP-ME in BS cell chloroplasts of WT (*m2m1*segregant – D; *m2m2* segregant – F), *Zmscr1-m2;Zmscr1h-m1* (E) and *Zmscr1-m2; Zmscr1h-m2*(G) leaves. Arrows in B, F and G indicate extra BS cells that are not in contact with the vasculature. Scale bars = 100μm.

Although double mutants did not produce deficiencies in the layer of BS cells around each vein, instances of supernumerary BS-like cells outside of the normal layer were observed (Fig. 5A, B). Quantification of this phenotype revealed significantly higher frequencies in both *Zmscr1;Zmscr1h* double mutants than in single mutants (Fig. 5C, Fig. 2K). However, even in the double mutants, over 90% of leaf veins had normal BS cell layers. To resolve whether the supernumerary BS-like cells were functionally equivalent to true BS cells, immunolocalization using an antibody against NADP-malic enzyme (NADP-ME) was carried out with wild-type and double mutant leaves. NADP-ME accumulated specifically in BS chloroplasts of wild-type maize leaves (Fig. 5D, E) and was detected in all of the supernumerary BS-like cells in double mutant leaves (Fig. 5F, G). Given the BS identity, cell position, and low frequency occurrence of these supernumerary cells, it is most likely that they are formed by late divisions of cells that are already differentiated as BS in the layer around the vein.

### *ZmSCR1* and *ZmSCR1h* transcripts accumulate in ground meristem cells of leaf primordia

As the protein sequences of both ZmSCR1 and ZmSCR1h predict cell-autonomous function, we sought to determine where the genes might act by determining spatial and temporal patterns of transcript accumulation during early leaf development. To this end, *in situ* hybridization was undertaken with developing wild-type leaf primordia. Fragments in the first exon (*ZmSCR1*) or the 3’UTR (*ZmSCR1h*), that were predicted to be gene-specific (Fig. S3), were used to distinguish *ZmSCR1* and *ZmSCR1h*. In plastochron (P)4 and P5 leaf primordia, transcripts of both genes were detected in the layer of ground meristem cells that surrounds developing veins, but were not detected in dividing procambial centres or in procambial-derived BS precursor cells (Fig. 6A-H). Notably, high transcript levels were observed in the single M precursor cells that are present between developing veins at P4 (Fig. 6B, C, E-H). These cells divide to form the two M cells present in the recurring vein-BS-M-M-BS-vein units that characterise Kranz anatomy in maize. Transcripts could not be detected in the regions of primordia where 2 interveinal cells were already present (Fig. 6G, H). The observed patterns of gene expression led us to hypothesise that ZmSCR1 and ZmSCR1 h pattern the cell layer encircling the BS during the development of Kranz anatomy.

**Figure 6.**
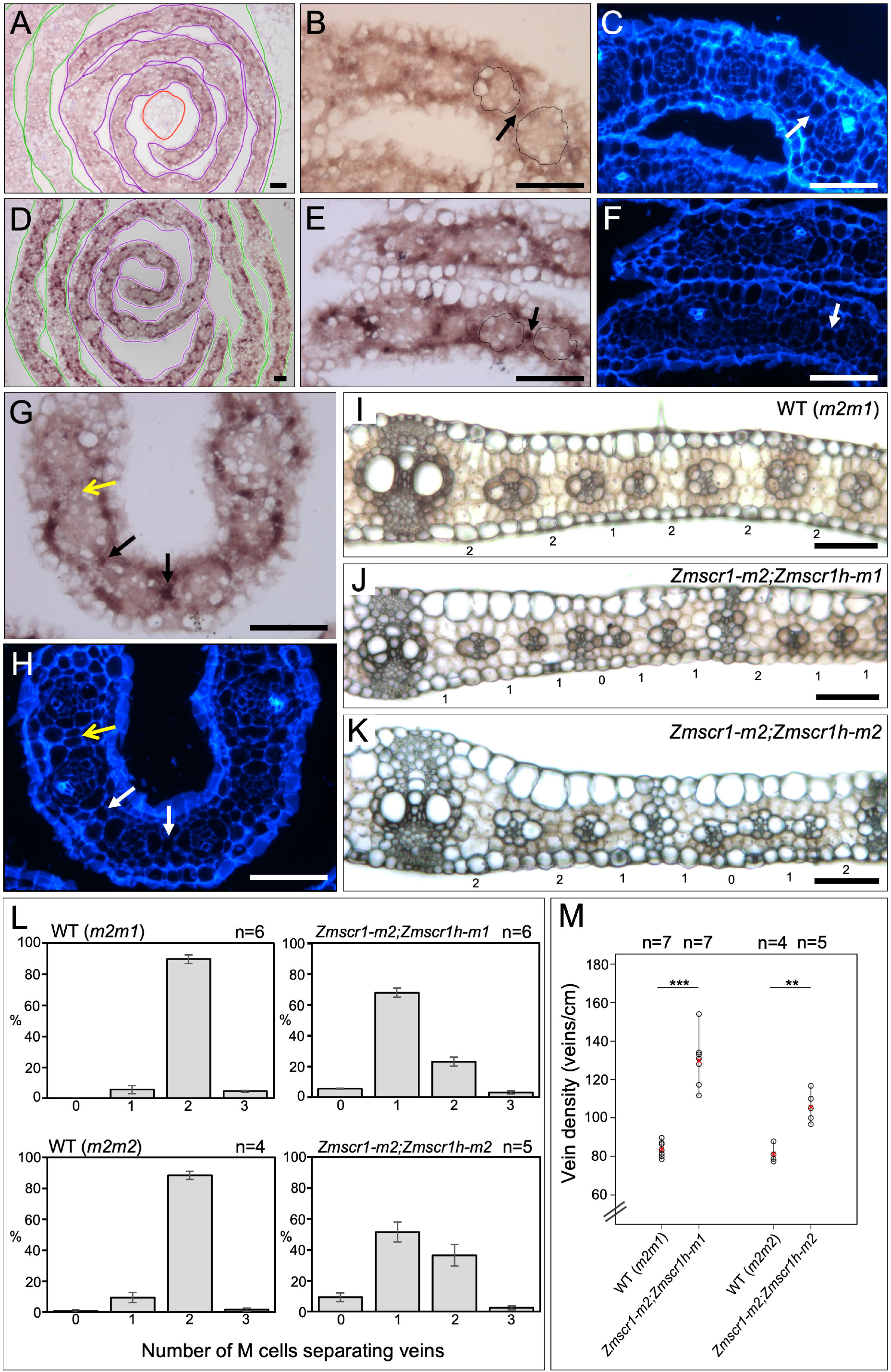
ZmSCR1 and ZmSCR1h regulate divisions of M cell precursors. **A-H)** *In situ* hybridization of *ZmSCR1* (A-C) and *ZmSCR1h* (D-H) in developing leaf primordia. Plastochron (P) numbers are indicated by coloured outlines: red = P3; purple = P4; green = P5 (A and D). Higher magnification P4 cross sections were imaged under both brightfield (B, E, G) and UV (C, F, H) illumination to show hybridization signal and calcofluor staining of cell walls respectively. Closed black and white arrows indicate single M precursor cells separating developing veins, open yellow arrows indicate two M precursor cells separating developing veins. Scale bars = 50μm (A-H). **I-K)** Representative fresh transverse sections of WT (*m2m1* segregant) (I), *Zmscr1-m2;Zmscr1h-m1* (J) and *Zmscr1-m2;Zmscr1h-m2* (K) fully expanded leaf 5. Scale bars = 100μm. Numbers below each section are the number of M cells separating the vascular bundles. Note that (I) is the same image as Fig. 5A. **L)** Quantification of the mean % of vascular bundles separated by 0-3 M cells in WT and *Zmscr1;Zmscr1h* fully expanded leaf 5. Error bars are standard error of the mean. **M)** Quantification of vein density in *Zmscr1;Zmscr1h* double mutants and corresponding WT segregants. Means are indicated by red crosses. Statistical significance between WT and mutants was assessed using Student’s t-tests (two-tailed): ** indicates p ≤ 0.01; *** indicates p ≤ 0.001. In (L) and (M) WT (*m2m1*)is also presented in Fig. 2I&J.

### Impaired mesophyll cell divisions are associated with higher vein density in *Zmscr1;Zmscr1h* mutant leaves

Consistent with transcript accumulation in M precursor cells between developing veins of WT P4 leaf primordia, there was a marked increase in the number of veins separated by just a single M cell in mature fully expanded leaves of *Zmscr1;Zmscr1h* mutants (Fig. 6I-L). In segregating wild-type backgrounds (and in single mutants – Fig. 2I), around 90% of all veins were separated by two M cells; whereas in both *Zmscr1-m2;Zmscr1h-m1* and *Zmscr1-m2;Zmscr1h-m2* mutants, most veins were separated by one M cell (68% and 52% of veins surveyed respectively) (Fig. 6L). Veins with no M cells in between were observed at 5-10% frequency in double mutants. This low frequency phenotype could result from the failure to specify a M precursor cell (and any subsequent divisions) or from ectopic BS cell divisions displacing M cells, either of which would effectively lead to vein anastomoses. The more penetrant phenotype suggests that ZmSCR1 and ZmSCR1h act redundantly to promote the cell division that generates two M cells in the characteristic vein-BS-M-M-BS-vein unit of Kranz.

To determine whether the decrease in M cell number between veins resulted in higher vein density across the leaf, or whether a compensatory reduction is vein number was manifest, vein number and leaf width were quantified in wild-type and double mutant leaves. Figure 6M shows that total vein density is significantly increased in *Zmscr1;Zmscr1h* double mutants. As well as reflecting fewer M cell divisions, this phenotype could also reflect the reduced width of *Zmscr1;Zmscr1h* mutant leaves (Fig. S2B), as both leaf width and cell size influence vein density in monocots. Either way, the increased vein density in *Zmscr1;Zmscr1h* mutants does not reflect an increase in the total number of veins formed across the width of the leaf (Fig. S2E), and as such a direct role for ZmSCR1 and ZmSCR1 h in the initiation and development of leaf veins can be eliminated.

### The relative proportions of intermediate vein ranks is altered in *Zmscr1;Zmscr1h* mutants

Although a role for *ZmSCR1* and *ZmSCR1h* in regulating the number of leaf veins was discarded, quantification of vein numbers and ranks in *Zmscr1;Zmscr1h* mutants revealed a striking shift in the proportion of each vein type that developed. *Zmscr1;Zmscr1h* mutants formed lignified sclerenchyma ad- and abaxially to the vein at far greater frequency than in wild-type (Fig. 7). In both C3 and C4 monocots, sclerenchyma is associated with both lateral and rank-1 intermediate veins. Rank-2 intermediate veins that do not form lignified sclerenchyma only form in C4 leaves with Kranz anatomy. In a typical wild-type maize leaf, two lateral veins are separated by between one and three evenly spaced rank-1 intermediate veins. These are in turn separated by multiple rank-2 intermediate veins, leading to a ratio of rank-1 to rank-2 veins of less than 0.2 (indicating on average >5 rank-2 veins to every rank-1 vein) (Fig. 7A, C, E). In contrast, a striking and consistent ratio of around 0.5 (indicating only two rank-2 veins to every rank-1 vein) is observed in *Zmscr1;Zmscr1h* double mutants (Fig. 7B, D, E), a ratio that is much greater than that seen in single *Zmscr1-m2* mutants (Fig. 2J). Notably, the extra rank-1 intermediates are not evenly spaced as in wild-type, but instead can be immediately adjacent to other rank-1 intermediate veins (Fig. 7D), and sclerenchyma is preferentially positioned on the abaxial side of the vein. Given that *ZmSCR1* and *ZmSCR1h* transcripts accumulate in ground meristem cells both ab- and adaxially to developing veins (Fig. 6 A-H), the observed shift in vein ranks in double mutants might suggest that in wild-type leaves ZmSCR1/ZmSCR1 h promote cell divisions and/or specification of M cells in these regions, and in so doing suppress sclerenchyma formation.

**Figure 7.**
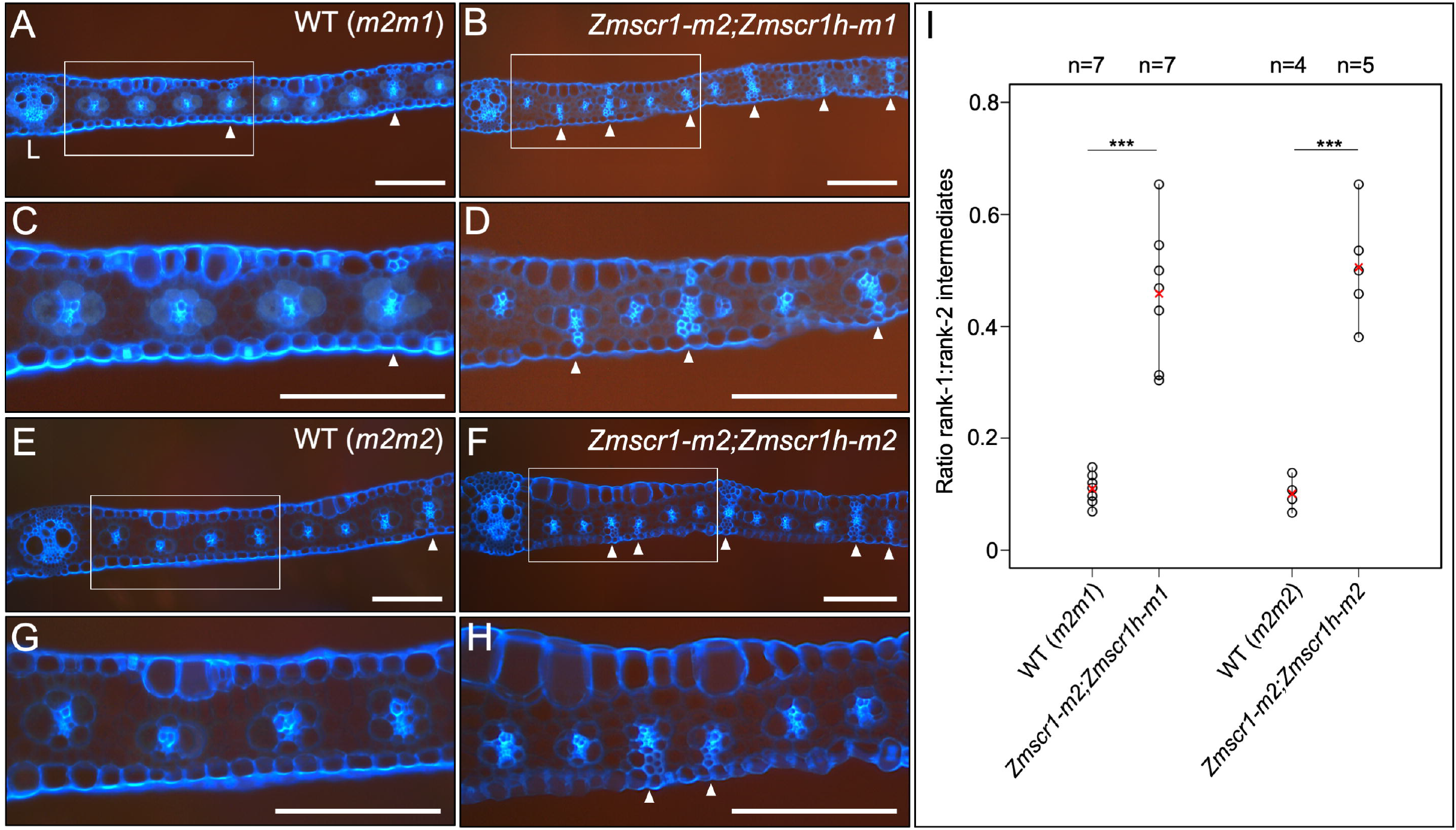
*Zmscr1;Zmscr1h* mutants form altered ratios of rank-1 to rank-2 intermediate veins. **A-H)** Representative fresh cut transverse sections of fully expanded leaf 5 from WT (*m2m1* segregant – A, C; *m2m2* segregant – E, G) and mutant (*Zmscr1-m2;Zmscr1h-m1* – B, D; *Zmscr1-m2;Zmscr1h-m1* – F, H) leaves imaged under UV illumination to show fluorescence associated with lignin. White boxes in A, B, E and F indicate an area that is enlarged in C, D, G and H respectively. White arrows indicate rank-1 intermediate veins with accompanying lignified sclerenchyma. L indicates a lateral vein. Scale bars = 100μm. **E)** Quantification of the ratio of rank-1 to rank-2 intermediate veins. Means are indicated by red crosses. Statistical significance between WT and mutants was assessed using Student’s t-tests (two-tailed): *** indicates p ≤ 0.001. WT (*m2m1*) data is also presented in Fig. 2J.

## DISCUSSION

Pattern formation is a fundamental process in both plant and animal development. In plants, radial patterning around the vasculature is of particular importance, and in the root is regulated by the SHR/SCR pathway (Cui et al., 2007). Here, we have shown that maize encodes two *SCR* genes which are equally orthologous to *AtSCR* (Fig. 1), and that *Zmscr1;Zmscr1h* double mutants exhibit a perturbed growth phenotype (Fig. 3) which closely resembles that seen in *Atscr* mutants (Di Laurenzio et al., 1996). The roots of *Zmscr1;Zmscr1h* plants lack an organized endodermal cell layer (Fig. 4), consistent with the reported localization of *ZmSCR1* transcripts in the developing endodermis (Lim et al., 2005). However, this phenotype is somewhat distinct from *Atscr* mutants in which a single organized ground-tissue layer with features of both the endodermis and cortex surrounds the vasculature (Di Laurenzio et al., 1996). In leaves, *ZmSCR1* and *ZmSCR1h* transcripts accumulate preferentially in ground meristem cells that will divide and differentiate into M cells (Fig. 6A-H), and fewer M cells are found in mature leaves of *Zmscr1;Zmscr1h* mutants than in wild-type (Fig. 6I-L). All of the phenotypic perturbations observed in double mutants are either absent or significantly less frequent in single mutants (Fig. 2), indicating that ZmSCR1 and ZmSCR1h function redundantly. Collectively, these results demonstrate that the SCR radial patterning mechanism operates in both roots and leaves of maize.

The canonical SHR/SCR pathway, in which AtSCR prevents AtSHR movement more than one cell-layer away from the root vasculature, was characterized in the context of roots with a single layer of both endodermis and cortex (Cui et al., 2007). However, maize and other monocots form multiple cortex layers (Clark and Harris, 1981; Dolan et al., 1993; Hochholdinger et al., 2018; Wu et al., 2014). It has been proposed that the number of cortex cell-layers in monocots is regulated by the extent to which SHR moves away from the root vasculature (Henry et al., 2017; Wu et al., 2014). Although SHR movement has not been confirmed in maize, there is no obvious change in the number of cortex cell-layers formed in *Zmscr1;Zmscr1h* mutants, despite the endodermal layer being absent (Fig. 3). This can be explained in one of three ways; 1) SHR is not involved in regulating cortical cell-layers in maize; 2) SCR does not restrict SHR movement in maize roots, a suggestion supported by the finding that movement of monocot SHR proteins was not constrained in planta by interaction with AtSCR (Wu et al., 2014); or 3) SHR is necessary but not sufficient to induce the formation of extra cell-layers, consistent with the finding that *Atscr* mutants form only one ground-tissue layer despite SHR movement being unconstrained (Di Laurenzio et al., 1996). Crucially, all of these alternatives indicate that the canonical SHR/SCR pathway is modified in roots that develop multiple layers of cortex.

The root endodermis and leaf BS are considered analogous (Esau, 1943; Nelson, 2011), and as such it has been suggested that the differentiation of both cell-types is regulated by the same genetic mechanism (Slewinski, 2013). However, in *Zmscr1;Zmscr1h* mutants an organized root endodermis is absent, whereas all leaf veins have a ring of surrounding BS cells (Fig. 5). In some cases, supernumerary BS cells that resemble those seen in the maize *tangled1* mutant (Jankovsky et al., 2001), are also observed around leaf veins (Fig. 5). In the *tangled1* mutant these supernumerary cells result from abnormal late divisions caused by perturbations in cell-division planes throughout the leaf (Jankovsky et al., 2001), suggesting that similar compensatory cell divisions may occur in *Zmscr1;Zmscr1h* mutants in response to aberrant divisions of ground meristem cells. Although a role for ZmSCR1 and ZmSCR1h in specifying BS cell fate remains formally possible, given that there is no evidence of preferential BS expression either early in leaf development (Fig. 6A-H) or in mature leaves (Chang et al., 2012; Denton et al., 2017; Li et al., 2010; Tausta et al., 2014) and that amino acid sequences predict both proteins are immobile (Gallagher and Benfey, 2009), a role in M patterning is more likely. Perturbations in *Zmscr1;Zmscr1h* leaves validate this suggestion in that most veins are separated by only one M cell, indicating impaired division of the single M-precursor cell that is marked by high *ZmSCR1* and *ZmSCR1h* transcript accumulation (Fig. 6). In addition, sclerenchyma forms ab- and adaxially to veins where M cells would normally develop (Fig. 7), suggesting that ZmSCR1 and ZmSCR1h inhibit the longitudinal divisions that give rise to sclerenchyma (Bosabalidis et al., 1994; Esau, 1943) and/or promote M cell differentiation. Taken together, these results refute the hypothesis that the endodermis and BS are patterned by the same mechanism, and instead suggest that SCR functions to promote the development of endodermal cells in the root and M cells in the leaf.

The most consistent mutant phenotype in *Zmscr1;Zmscr1h* leaves is the increased number of veins separated by only one M cell, which accounts for 68% (*Zmscr1-m2;Zmscr1h-m1*) or 52% (*Zmscr1-m2;Zmscr1h-m2*) of veins in double mutants compared to <10% in WT (Fig. 6L). However, penetrance is clearly not complete as 23% and 36% of veins are still separated by two M cells (Fig. 6L). Notably, this contrasts with complete penetrance of the endodermal defects in *Zmscr1;Zmscr1h* mutant roots (Fig. 3). We hypothesize that this difference reflects single versus multiple clonal origins of endodermal and M cells. At least in Arabidopsis, all endodermal cells arise from divisions of initials that are distinct from those that form the vasculature (Dolan et al., 1993). By contrast, although the central ground meristem layer gives rise to all leaf veins and BS cells plus M cells in that layer (Langdale et al., 1989), once procambium has been specified two origins of central M cells can be distinguished (Jankovsky et al., 2001). Lineage analyses found that 67% of sectors induced after procambium initiation comprised a complete ring of BS cells but no M cells, whereas 33% of sectors consisted of a few cells in the BS layer plus one or more adjacent M cells (Jankovsky et al., 2001). As such, two thirds of M cells in the central leaf layer originate from ground tissue that is clonally distinct from the BS whereas one third are clonally related to adjacent BS cells. These proportions are consistent with the percentages of veins separated by one (~60%) versus two (~40%) M cells in *Zmscr1;Zmscr1h* mutants, and thus suggest that the division and differentiation of M cells originating from the same precursor cell as the vascular bundle is regulated by a SCR-independent mechanism. This implies that the transition from ground meristem to M cell is regulated by at least two distinct mechanisms within the maize leaf.

It has been hypothesized that SHR/SCR mediated patterning of cell-types in the leaf is specific to C4 Kranz anatomy (Slewinski, 2013). Current evidence is supportive of Kranz-specific SHR/SCR roles in that *Atscr* mutant leaves exhibit only a slight enlargement of BS cell size (Cui et al., 2014) and constitutive expression of *ZmSCR1* in rice failed to disrupt any aspect of leaf development (Wang et al., 2017). These observations suggests that SCR is neither necessary nor sufficient to regulate the spatial arrangement of cell-types in the inner leaf layers of C_3_ plants. Transcripts of one of the two rice *SCR* orthologs localize to cells that give rise to stomata (Kamiya et al., 2003) and constitutive expression of *ZmSHR2* in rice induces changes in stomatal rather than BS or M cell patterning (Schuler et al., 2018), suggesting that the SHR/SCR pathway may regulate cell-type specification in the epidermis rather than the inner leaf layers of rice. However, this suggestion needs further investigation given that our data reveal a role for SCR in M cell development and that M cells are a common feature of both C3 and C4 leaves. Based on current evidence we conclude that the SHR/SCR pathway represents a flexible regulatory module that has been co-opted to pattern cell-types in a range of developmental contexts in both roots and shoots of flowering plants.

## MATERIALS AND METHODS

### Plant stocks and growth conditions

UniformMu seed stocks harbouring *Mutator* insertions in either *ZmSCR1* (GRMZM2G131516) or *ZmSCR1h* (GRMZM2G015080) were acquired from the maize genetics COOP stock centre (http://maizecoop.cropsci.uiuc.edu) (Fig. S1A). Plants were grown in the field at Iowa State University, and individuals harbouring the *Zmscr1-m2* allele were outcrossed to those harbouring either the *Zmscr1h-m1* or *-m2* allele. F1 plants heterozygous for both insertions were self-pollinated to yield F2 populations segregating 1/16 for both segregating wild-type and *Zmscr1;Zmscr1h* double mutants. Self-pollinations of *Zmscr1-m2/+;Zmscr1h-m1* F2 plants resulted in F3 families where *Zmscr1-m2;Zmscr1h-m1* homozygous double mutants segregated 1/4. The inbred line B73 was used for *in situ* hybridization experiments.

For developmental analyses, plants were grown in a greenhouse in Oxford with a 16hr/8hr light regime. Daytime temperature was maintained at 28°C and night-time temperature at 20°C. Supplemental light was provided when natural light was lower than 120μmol photon m^-2^ s^-1^. Seed were germinated in warm, damp vermiculite and transferred after one week to 12cm diameter pots containing a 3:1 mix of John Innes No.3 Compost (J. Arthur Bower) and medium vermiculite (Sinclair Pro).

### Genotyping

A first round of genotyping was undertaken on genomic DNA extracted with a sodium dodecyl sulfate (SDS) high throughput 96-well plate protocol. Leaf tissue was homogenized with 500μl SDS extraction buffer (200mM Tris pH 7.5, 250mM NaCl, 25mM EDTA, 0.5% SDS) in 96-well collection microtubes. Plates were then centrifuged at 6000rpm for 10 min, and 200μl supernatant removed and mixed with 200μl isopropanol in a 96 well polypropylene plate. After 10 min incubation at room temperature, plates were centrifuged and the supernatant discarded. DNA pellets were washed in 70% (v/v) ethanol, centrifuged and air dried before being resuspended in 100μl dH_2_O.

Individuals identified with genotypes of interest were subjected to a second round of genotyping using genomic DNA extracted using a modified cetyl-trimethyl-ammonium bromide (CTAB) protocol optimized to yield high-quality DNA (Murray and Thompson, 1980). Leaf tissue was homogenized at room temperature in CTAB buffer (1.5% (w/v) CTAB, 75mM Tris-HCl pH 8, 15mM EDTA pH 8, 1.05M NaCl) and heated to 65°C for 30 min. An equal volume of 24:1 chloroform: isoamyl alcohol was added and mixed, before samples were centrifuged. The resultant supernatant was mixed with 2.5 volumes of 100% (v/v) ethanol. The precipitate was collected by centrifugation and washed with 70% (v/v) ethanol before drying and resuspending in 100μl dH_2_O.

The presence of mutant alleles and the sites of insertion were elucidated by PCR using a 1:1 mix of two primers (EOMu1 and EOMu2, Fig. S2B, C) designed to amplify out from both the 5’ and 3’ end of the *Mutator* element, and a primer amplifying from the gene sequence adjacent to the predicted insertion site (Fig. S2B, C). The presence of the wild-type allele was confirmed using a pair of primers that flanked the insertion site (Fig. S2B, C). In some cases, the ‘wild-type’ primers amplified across the transposon producing a larger product size from mutant alleles than from wild-type alleles. However, in most cases there was no amplification with these primers when only the mutant alleles were present. PCR amplifications were carried out in a total reaction volume of 10μL containing 5 μL of 2xGoTaq master mix (Promega) and 2.5 μL of 4M betaine. Reaction conditions were as follows: 95°C for 5 min; 35 cycles of 95°C for 30 s, 57-64°C for 30 s, and 72°C for 1:00-1:30 min; and 72°C for 10 min. All PCR experiments were designed and tested using homozygous single mutant lines to ensure that primers only amplified from the correct gene sequence. All PCR products were assessed by agarose gel electrophoresis.

To determine whether transposons were retained in the transcripts from mutant alleles, RNA was extracted using an RNeasy Plant Mini Kit following the manufacturer’s instructions (Qiagen). Extracted RNA was DNaseI treated using TURBO DNase (Invitrogen) and 2μg RNA was used for cDNA synthesis using a Maxima First Strand cDNA synthesis kit (Thermo Scientific). RT-PCR was carried out on 1/10 cDNA dilutions using primers amplifying from the transposon to the flanking genomic region (Fig. S2).

### Analysis of fresh leaf sections

Plants were photographed 32 days after planting, and fully expanded leaf 5 (i.e. the fifth leaf to emerge after germination) was removed at the ligule for phenotypic analysis. Leaf length, width and midrib extension (the point along the proximal/distal axis at which the midrib was no longer visible) were recorded. Segments of leaf encompassing the midrib and the 3-4 adjacent lateral veins were cut from the midpoint along the proximal/distal axis and positioned upright in 7% agar. Once cooled, blocks were trimmed and mounted such that veins were vertically orientated. 50-60μm sections were cut using a vibratome and then cleared for around 10 min in 3:1 ethanol: acetic acid. Sections were incubated in 70% ethanol overnight, then floated on slides with 70% ethanol (v/v) and covered with a coverslip. Leaf sections were imaged using a Leica DMRB microscope with QImaging MicroPublisher camera (QImaging, www.qimaging.com) and Image-Pro Insight software (MediaCybernetics, www.mediacy.com). Images were taken using brightfield (which enabled BS and M cells to be identified) and UV (which enabled sclerenchyma and thus vein orders to be determined) illumination. Images were tiled together so that the region of leaf between two lateral veins was represented. Subsequent quantification of segment width was undertaken using the ImageJ software package (www.imagej.nih.gov).

### Tissue fixation and embedding

Segments of primary and seminal roots were fixed in ice-cold 90% acetone for 15 min, rinsed with 100mM phosphate buffer (pH 7), placed in 3:1 ethanol: acetic acid for a further 15 min and then transferred to 70% (v/v) ethanol. B73 shoot apices were harvested on ice after 7 days growth, prior to the emergence of the first leaf through the coleoptile. Harvested apices were fixed and vacuum infiltrated for 1 min in ice-cold 4% (w/v) paraformaldehyde. Fresh paraformaldehyde was added following vacuum infiltration and samples left overnight. The following day, samples were dehydrated through ice cold 10%, 30%, 50% and 70% (all v/v) ethanol for 2 hours each. Segments of leaf were cut from the midpoint along the proximal/distal axis of fully expanded leaf 5, encompassing ~3 lateral veins adjacent to the midvein. Leaf segments were fixed for 30 min in 3:1 ethanol: acetic acid and transferred to 70% EtOH. All fixed tissue was stored at 4°C in 70% ethanol (v/v), and prior to embedding root samples were placed in 0.7% agar.

Fixed tissue was dehydrated and embedded in paraffin wax using a Tissue-Tek VIP machine (Sakura, www.sakura.eu). Samples were dehydrated at 35°C through 70%, 80%, 90% (with 1% (w/v) eosin), 95% and three times 100% (all v/v) ethanol for 1 hour each. Samples were then incubated at 35°C in 3 times histoclear for 1 hour each. Finally, samples were wax-infiltrated by four incubations of 2 hours in paraffin at 65°C. Embedded tissue was then placed in wax blocks and left to solidify at 4°C overnight. Wax blocks were trimmed and 10μm transverse sections cut using a Leica RM2135 rotary microtome and placed on slides at 37°C to dry overnight.

### Toluidine blue staining of root sections

Slides were placed in histoclear twice for 10 min each to remove wax, before being taken through an ethanol re-hydration series (1 min in each). Slides were stained in 0.05% (w/v) Toluidine blue (50mM citrate buffer, pH 4.4) for 5 seconds, rinsed in dH_2_O and then dried and mounted using a drop of entellen (Merck Millipore). Images were taken using brightfield illumination as above.

### *In situ* hybridization

*In situ* hybridization was carried out using wax-embedded shoot apices as described by Schuler et al., 2018, with digoxygenin (DIG)-labelled RNA probes designed to specifically detect either *ZmSCR1* or *ZmSCR1h* transcripts (Fig. S3). The *ZmSCR1* probe was a 108bp region towards the end of the first exon, which shared 78% identity with the corresponding region of *ZmSCR1h*. The *ZmSCR1h* probe was a different 108bp region in the 3’UTR. The current predicted gene-model for *ZmSCR1* does not include the majority of this 3’UTR region (Phytozome12), and as such the probe should be highly specific for *ZmSCR1h*. If this gene-model is incorrect, and *ZmSCR1* encodes a longer 3’UTR, then the *ZmSCR1h* probe shares 63% sequence identity with the *ZmSCR1* gene (Fig. S3). Post-hybridization washes were undertaken with 0.005x (*ZmSCR1*) and 0.01x (*ZmSCR1h*) SSC buffer made from a 20x SSC stock (3M NaCl, 0.3M Na3citrate), calculated to ensure stringency.

### Immunolocalization

Slides were dewaxed twice for 10 min in histoclear, then transferred through 99% (x2), 95% and 85% ethanol (all v/v) for 2 min each. Slides were incubated in 3% (v/v) H_2_O2 in methanol for 15 min, and rehydrated through 70%, 50% and 30% ethanol (all v/v) and finally dH_2_O twice. Slides were then incubated in PBS/BSA buffer (0.15M NaCl, 10mM phosphate buffer pH 7, 1mgml^-1^ BSA) for 5 min, drained and incubated for 15 min in 0.1% (w/v) Goat IgG (Sigma Aldrich) (in PBS/BSA) and rinsed in PBS/BSA. Slides were then incubated for 15 min with a 1/1000 dilution (in PBS/BSA) of ZmNADP-ME antibody (Langdale et al., 1987), before being rinsed twice for 15 min each in PBS/BSA and incubated for 15 min in a 1/100 dilution (in PBS/BSA) of biotinylated goat anti-rabbit secondary antibody (Sigma Aldrich). Slides were rinsed as before and incubated with a 1/100 dilution (in PBS/BSA) of biotinylated/streptavidin/horseradish peroxidase complex (GE Healthcare) before a final PBS/BSA rinse. SIGMAFAST™ 3,3’-Diaminobenzidine (DAB) tablets (Sigma Aldrich) were used to prepare a staining solution as per the manufacturer’s instructions including 0.03% (w/v) NiCl2. Slides were covered in staining solution and observed until colour had developed sufficiently (usually ~1 min), before being rinsed in dH_2_O and dehydrated through the original ethanol series. Finally, slides were mounted using DPX (Sigma Aldrich) and visualized using brightfield microscopy in the same way described for leaf and root histology.

### Quantification and statistics

Quantification of leaf segment width was undertaken using the ImageJ software package (www.imagej.nih.gov). Statistical analysis was undertaken using RStudio (www.rstudio.com). Student’s t-tests were used to test for differences in leaf length, leaf width, midvein extension, vein density and vein order ratios between mutants and the corresponding segregating WT lines. Standard errors of the mean were calculated for M and BS data.

### Phylogeny construction

Primary transcript proteomes from eleven species were downloaded from Phytozome12 (Goodstein et al., 2012); *Zea mays* (B73), *Sorghum bicolor, Setaria italica, Setaria viridis, Oryza sativa, Brachypodium distachyon and stacei*, and *Ananas comosus* were chosen for monocots, *Arabidopsis thaliana* and *Solanum lycopersicum* for dicots and *Physcomitrella patens* as an outgroup. The ZmSCR1 (GRMZM2G131516) primary protein sequence was used as a query in a BLASTp search (evalue of 1e-3) against this proteome database, with the top 100 hits retained. The gene model from one sorghum *SCR* ortholog (Sobic008G023401.1) was incorrect, as it was predicted to begin without a start codon. The upstream region of this sequence was interrogated, and an in-frame start codon was identified. No suitable expression data were available to validate this corrected gene model, but as the original model was definitely incorrect, the new version was used for alignment. The 100 sequences were aligned using MergeAlign (Collingridge and Kelly, 2012), and the resultant alignment (File S1) was used to generate a maximum likelihood phylogeny using IQtree (Hoang et al., 2018; Trifinopoulos et al., 2016). In parallel, the alignment was trimmed using trimAl to remove poorly aligned regions, such that columns with less than 30% of the sequences represented were discounted from further analysis (Capella-Gutiérrez et al., 2009). Trimming did not alter the topology of the resultant tree, and as such the untrimmed version is presented here. Trees were visualized using the Interactive Tree of Life (iTOL) software (Letunic and Bork, 2016).

## ACKNOWLEDGEMENTS

We thank the UniformMu project and the Maize Genetics Stock Centre for providing the original seed stocks; Daniela Vlad and Julia Lambret Frotte for helpful discussions throughout the work and during the course of manuscript preparation; Erik Vollbrecht, Rubén Rellán-Álvarez and Ruairidh Sawers for help with crossing and fieldwork; John Baker for plant photography and Julie Bull for technical support. TEH and JAL wrote the manuscript with all authors contributing to the final draft.

## AUTHOR CONTRIBUTIONS

TEH carried out genotyping, phenotypic characterization and data analysis; OVS undertook in situ hybridization; HW and PWB assisted with crossing and fieldwork support; JAL conceived the study and provided funding.

---

**Supplemental Figure 1.**
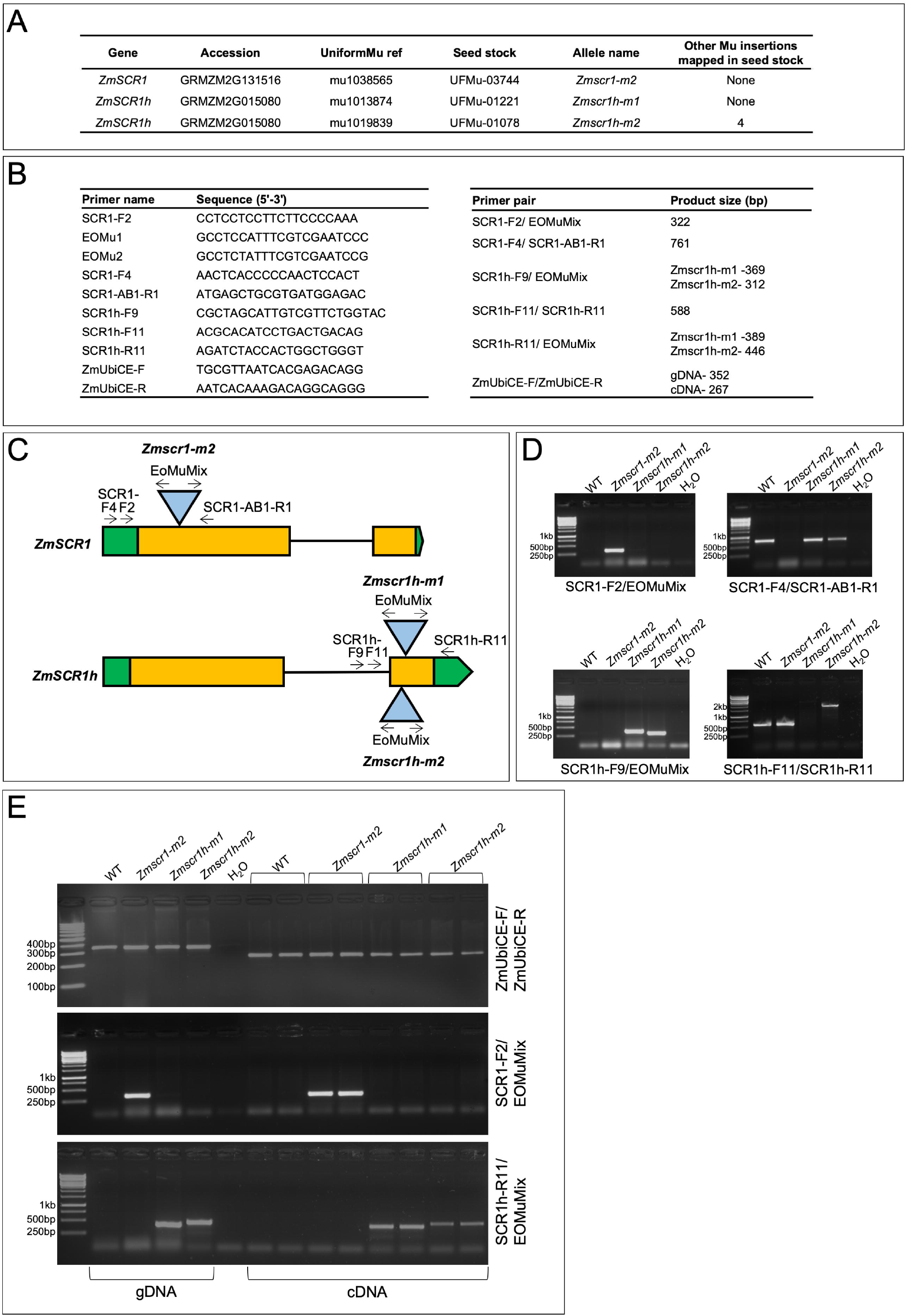
Validation of insertion alleles. **A)** Details of alleles provided by the UniformMu project. **B)** Primers used in this study. **C)** Schematic representation of each mutant allele, with the position of genotyping primers indicated. **D)** Agarose gels showing fragments amplified from WT and mutant alleles of both *ZmSCR1* and *ZmSCR1h*, with different primer pairs as indicated. Genomic DNA was used as template in all cases. **E)** Agarose gels of fragments amplified from genomic DNA or cDNA (copy of RNA extracted from leaf primordia), using primer pairs as indicated. Transposon sequences are present in the transcripts of each mutant allele.

**Supplemental Figure 2.**
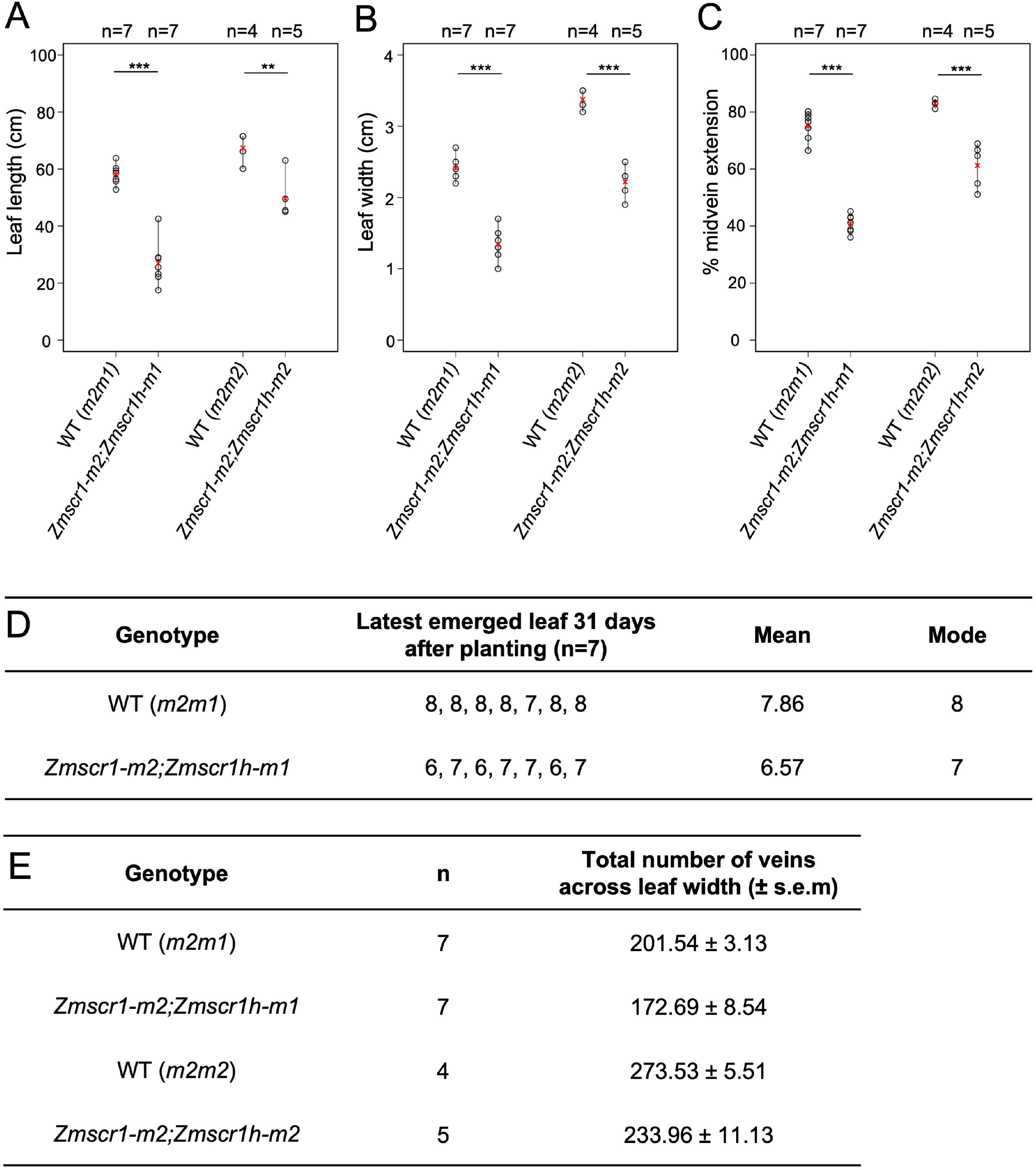
Quantification of *Zmscr1;Zmscr1h* growth characteristics. **A-C)** Quantification of leaf blade length from ligule to tip (A); leaf width at the midpoint along the proximal/distal axis (B); and the % extension of the midvein (C). Quantification was undertaken either 31 (*m2m1*) or 32 (m2m2) days after planting. Means are indicated by red crosses. Statistical significance between WT and mutants was assessed using Students t-tests (two-tailed): ** indicates p ≤ 0.01; *** indicates p ≤ 0.001. **D)** Quantification of the latest emerged leaf in segregating WT and *Zmscr1-m2;Zmscr1h-m1* mutants. **E)** Quantification of the total number of veins across the width of the leaf, calculated by multiplying leaf width by vein density. s.e.m indicates standard error of the mean.

**Supplemental Figure 3.**
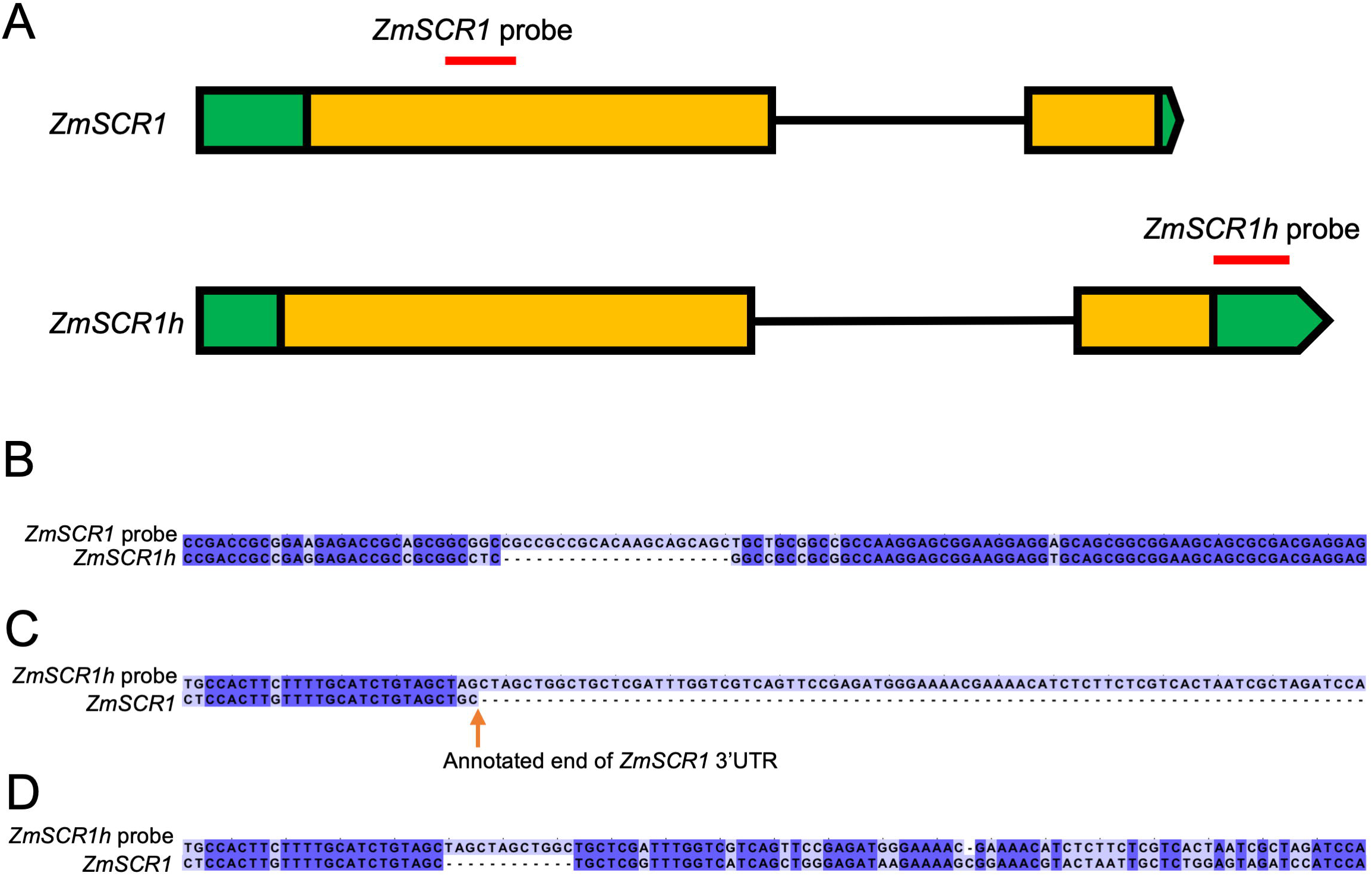
*In situ* hybridization probe design. **A)** Cartoon showing position of *ZmSCR1* and *ZmSCR1h* probes. Green indicates UTRs; orange indicates exons; black lines indicate introns; Red lines indicate hybridization probes. **B-D)** Alignment of the *ZmSCR1* probe with the *ZmSCR1h* gene sequence (B); the *ZmSCR1h* probe with the *ZmSCR1* annotated gene sequence (C); or the *ZmSCR1h* probe with the annotated *ZmSCR1* gene sequence and region downstream of the of the 3’UTR end (D). Dark blue indicates identical sequence, light blue indicates differences.

